# Fe^2+^/H_2_O_2_-mediated oxidation of homogentisic acid indicates the production of ochronotic and non-ochronotic pigments. Implications in Alkaptonuria and beyond

**DOI:** 10.1101/223099

**Authors:** Koen P. Vercruysse, Adam M. Taylor, Juan Knight

## Abstract

Homogentisic acid (HGA) can be oxidized by a combination of Fe^2+^ and H_2_O_2_ into a darkly colored high molecular mass pigment. Increasing the concentration of H_2_O_2_ can lead to the formation of a high molecular mass material that exhibits less absorbance in the visible range of the electromagnetic spectrum, while maintaining a strong absorbance in the UV range of the electromagnetic spectrum. FT-IR spectroscopy did indicate the presence of a chemical feature in the pigments generated through H_2_O_2_-mediated oxidation that is absent in pigments generated through air-mediated oxidation. Our observations could have implications in the pathophysiology of alkaptonuria. In alkaptonuria, patients suffer from homogentisic aciduria from birth, but develop ochronosis (darkening) of collagenous tissues much later in life due to the formation of a melanin-like pigment. Three major questions regarding ochronosis remain to be answered: 1) how is the pigment formed, 2) why does it appear by the third decade of life and 3) why is it sporadic in appearance? Our results suggest that ochronotic or non-ochronotic pigments can be generated from HGA depending on the oxidation reaction conditions. Thus, the absence of any visible pigment, as for younger alkaptonuria patients, could not necessarily mean the absence of HGA-derived melanin-like molecules. We compare our results and their potential implications for ochronosis to the changes in pigmentation observed in vitiligo or aging (greying) hair.

## 1. Introduction

Alkaptonuria (AKU) is a genetically-determined metabolic disease whereby the gene encoding the homogentisate 1,2 dioxygenase enzyme is mutated.[1, 2] Thus, AKU patients accumulate homogentisic acid (HGA) from birth. Around the third decade of life, AKU patients develop dark pigmentation in collagen-rich tissues, a condition termed ochronosis. The build-up of this ochronotic pigment leads to osteoarthropathy, but other tissues, e.g. cardiac valves, may deteriorate.[3-5] Although the biochemistry of the tyrosine metabolization pathway in normal and AKU patients is well established, three main questions regarding ochronosis remain: 1) how is the pigment synthesized, 2) why does it appear only in a later stage of life and 3) why is it sporadic, rather than blanket in its appearance?[6]

Using air-oxidation, HGA is readily converted into a darkly-colored, melanin (MN)-like pigment, but strong alkaline conditions appear to be required to initiate the process. [7] *In* vivo such conditions are unlikely to be the norm, however some cartilage samples display a neutral or alkaline pH. [8] A more likely means of oxygen-mediated oxidation and polymerization of HGA would require the presence of a catalyst; enzyme or other. Radical oxygen species (ROS) can serve as a source of oxidizing potential, particularly if transition metal ions like Fe^2+^ or Cu^2^+ are present. ROS-mediated oxidation is frequently simulated in laboratory settings by combining Fe^2+^ or Cu^2+^ with H_2_O_2_; the so-called Fenton chemistry. We have studied the oxidation of HGA using Fenton chemistry and evaluated the formation of MN-like pigment similarly as was done for the air-mediated oxidation of HGA under alkaline conditions.[7] We observed that, depending on the concentration of H_2_O_2_ used, a high molecular mass pigment material was generated with different absorbance properties in the visible range of the electromagnetic spectrum. Our experiments raise a hypothesis that a lesser-visible MN-like pigment can be generated from HGA depending on the intensity of the oxidizing conditions of the environment. In a broader context, our observations could be related to the depigmentation observed in vitiligo patients or in the graying of hair; for both of which increased levels of H_2_O_2_ has been implicated as a potential cause of the depigmentation.[9, 10]

## 2. Materials and Methods

### 2.1. Materials

Homogentisic acid was obtained from Sigma-Aldrich (St Louis, MO). FeCl_2_.2H_2_O was obtained from Fisher Scientific. H_2_O_2_ solution at 3% (v/v) was obtained from Kroger Co (Cincinnati, OH).

### 2.2. Small scale experiments

Mixtures containing various concentrations of HGA, Fe^2+^ and H_2_O_2_ were prepared in wells of a 96-well microplate and kept at room temperature (RT) for multiple days. Occasionally, photographs and absorbance readings at wavelengths between 400 and 600nm were taken. Details on concentrations involved in these reactions are provided in the Results section.

### 2.3. Large scale experiments

Reaction mixtures (total volume 25mL) containing about 100mg HGA, 0.3mM Fe^2+^ and 0.12% v/v (reaction A) or 1.2% v/v (reaction B) H_2_O_2_ were kept at RT for one week. Occasionally, aliquots from the mixtures were diluted with SEC solvent, centrifuged and analyzed using SEC. When the reactions were deemed complete, the mixtures were dialyzed and freeze-dried as described previously leading to pigments A and B.[7]

### 2.4. UV_Vis spectroscopy

UV/Vis spectroscopic measurements were made in wells of a 96-well microplate using the SynergyHT microplate reader from Biotek (Winooski, VT).

### 2.5. Size exclusion chromatography (SEC)

SEC analyses were performed as described previously.[7]

### 2.6. FT-IR spectroscopy

FT-IR spectroscopic analyses were performed as described previously.[7]

## 3. Results

### 3.1. Small scale experiments

Mixtures containing a fixed amount of HGA (5.5mM) and varying concentrations of Fe^2+^ (between 0 and 3mM) and H_2_O_2_ (between 0 and 0.2% v/v) were kept at RT. In the absence of any Fe^2+^ or H_2_O_2_, no changes in color were observed. In the presence of H_2_O_2_, between 0.01 and 0.2% v/v, dark colors were observed in the mixtures immediately upon the addition of H_2_O_2_. The intensity of the dark color increased with increasing concentrations of Fe^2+^. However, the mixtures containing 0.1 or 0.2% v/v H_2_O_2_ rapidly lost their initial dark color, to the point that some mixtures appeared colorless after overnight reaction. Fig. 1 shows a photograph of this experimental set up after two days of reaction.

**Fig.1:**
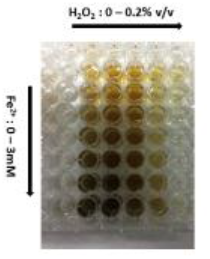
Photograph of mixtures containing a fixed amount of HGA (5.5mM) and varying concentrations of Fe^2+^ (between 0 and 3mM) and H_2_O_2_ (between 0 and 0.2% v/v) after two days of reaction at room temperature.

Fig. 2 presents the absorbance measured at 400nm as a function of H_2_O_2_ concentration for select reaction mixtures after two days of reaction at room temperature.

**Fig.2:**
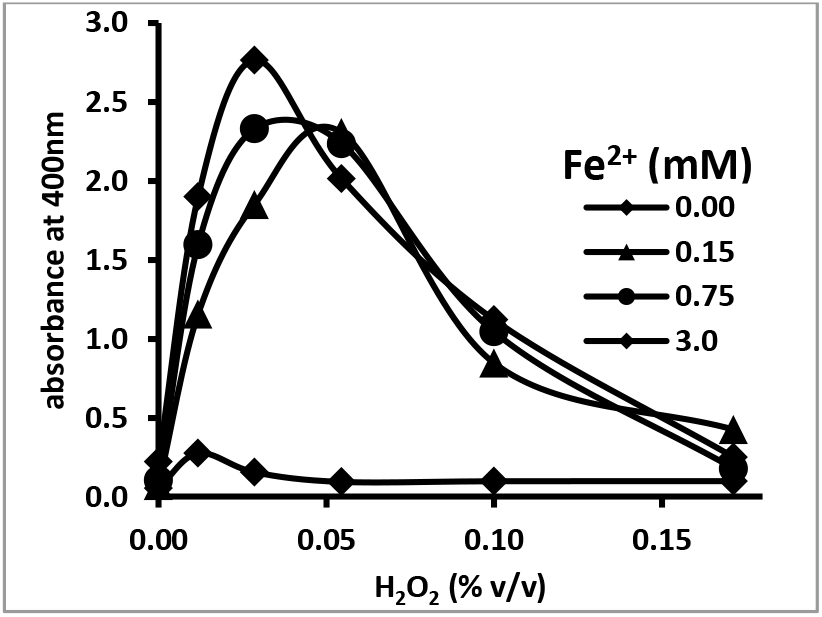
Absorbance at 400nm of reaction mixtures containing 5.5mM HGA and various concentrations of Fe^2+^ as a function of H_2_O_2_ concentration after two days of reaction at RT.

An aliquot of the almost colorless reaction mixture containing, 5.5mM HGA, 0.16mM Fe^2+^ and 0.1% H_2_O_2_ was diluted and analyzed by SEC. This analysis revealed the presence of a high molecular mass material, with significant absorbance in the UV range, but not in the visible range of the electromagnetic spectrum (results not shown). Mixtures containing a fixed amount of HGA (5.5mM) and varying concentrations of Cu^2+^ (between 0 and 3mM) and H_2_O_2_ (between 0 and 0.2% v/v) were kept at RT. A similar pattern of results as for the experiments involving Fe^2+^ were obtained. However, the reactions in the presence of Cu^2^+ appeared to proceed slower compared to the reactions in the presence of Fe^2+^ (results not shown). Mixtures containing a fixed amount of Fe^2+^ (0.1mM) and varying concentrations of HGA (between 0 and 20mM) and H_2_O_2_ (between 0 and 0.2% v/v) were kept at RT. Fig. 3 presents the absorbance measured at 500nm as a function of H_2_O_2_ concentration for select reaction mixtures after two days of reaction at room temperature.

**Fig.3:**
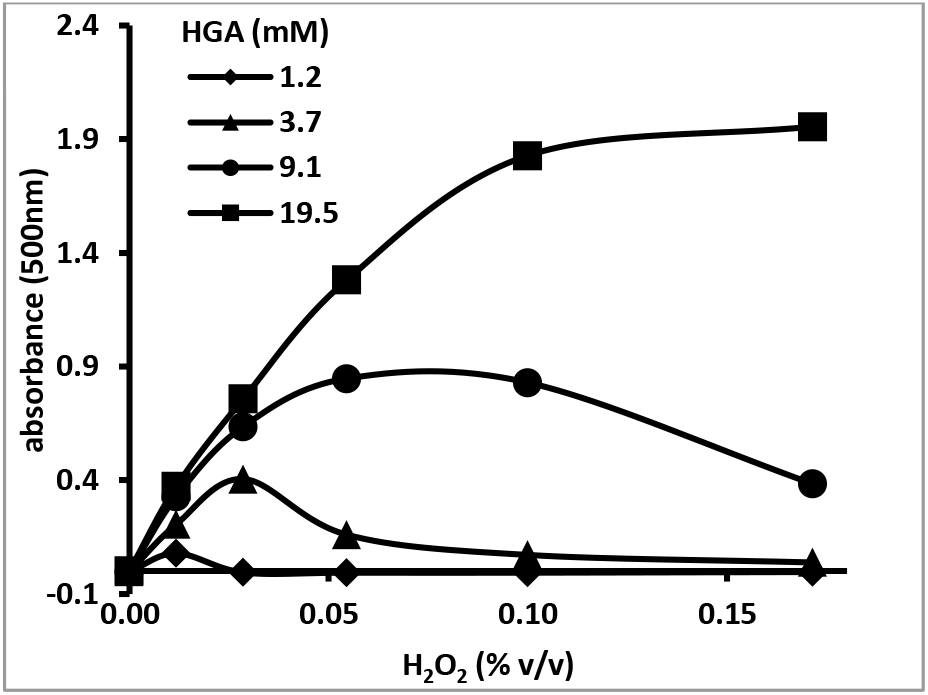
Absorbance at 500nm of reaction mixtures containing 0.1mM Fe^2+^ and various concentrations of HGA as a function of H_2_O_2_ concentration after two days of reaction at RT.

### 3.2. Large scale experiments

Upon the addition of H_2_O_2_ to reaction mixture A, the solution instantly turned black and remained as such throughout the remainder of the experiment. Upon the addition of H_2_O_2_ to reaction mixture B, the solution instantly turned yellow-brown and turned black within hours. After overnight standing, reaction mixture B had regained the yellow-brown color and remained as such throughout the remainder of the experiment. SEC analyses of the dialyzed reaction mixtures revealed a peak with retention times of 12.7 minutes (mixture A) and 12.5 minutes (mixture B) and no signs of peaks corresponding to HGA or H_2_O_2_ (results not shown). These retention times are similar to the SEC retention time, 12.4 minutes, of the pigment generated from HGA through air-oxidation in the presence of 1M NaOH. [7] The purified and dried pigment A consisted of a black, fibrous material, while the purified and dried pigment B material consisted of a brown, fibrous material. Both pigments were redissolved in water to a concentration of 0.2 mg/mL. Fig. 4 presents a photograph of these solutions and Fig. 5 presents the UV_Vis spectra recorded for both pigments.

**Fig.4:**
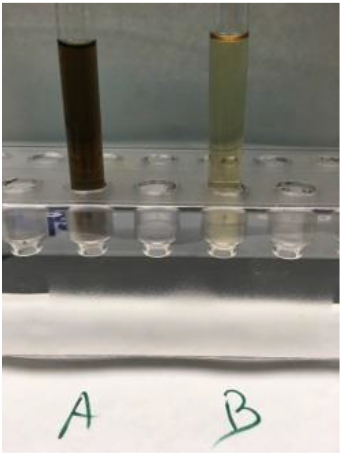
Photograph of solutions of purified pigment A (left) and pigment B (right) redissolved in water at 0.2 mg/mL.

**Fig.5:**
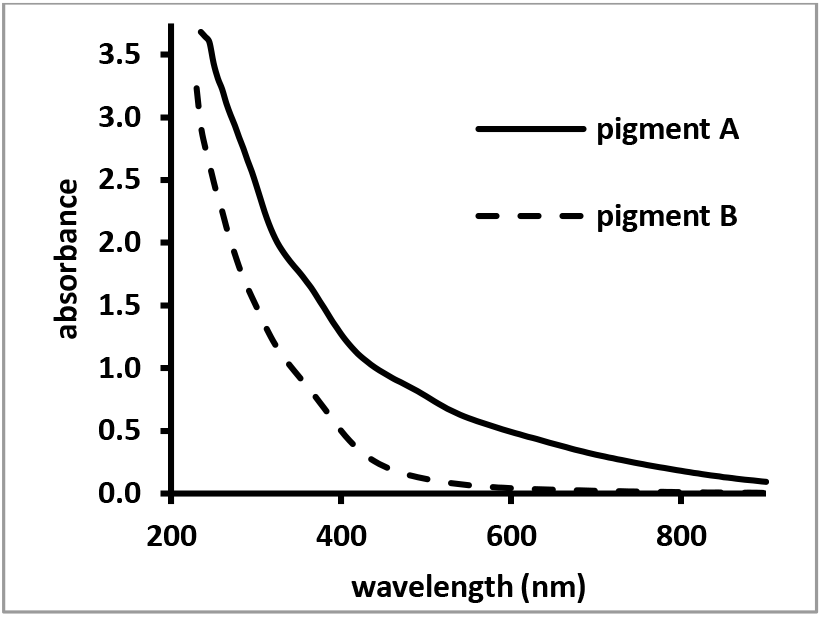
UV_Vis spectra of pigments A and B, prepared as discussed in this report, in water at 0.2mg/mL.

Comparing the UV_Vis spectra of pigments A and B, the absorbance in the Vis region of the electromagnetic spectrum of pigment B was much lower compared to pigment A. Both pigments still exhibited a strong absorbance in the UV range of the electromagnetic spectrum. Fig. 6 compares the FT-IR spectra of pigments A and B to the spectrum of a freshly prepared pigment derived from HGA through air oxidation in the presence of 1M NaOH.(pigment #5 in [7]

**Fig.6:**
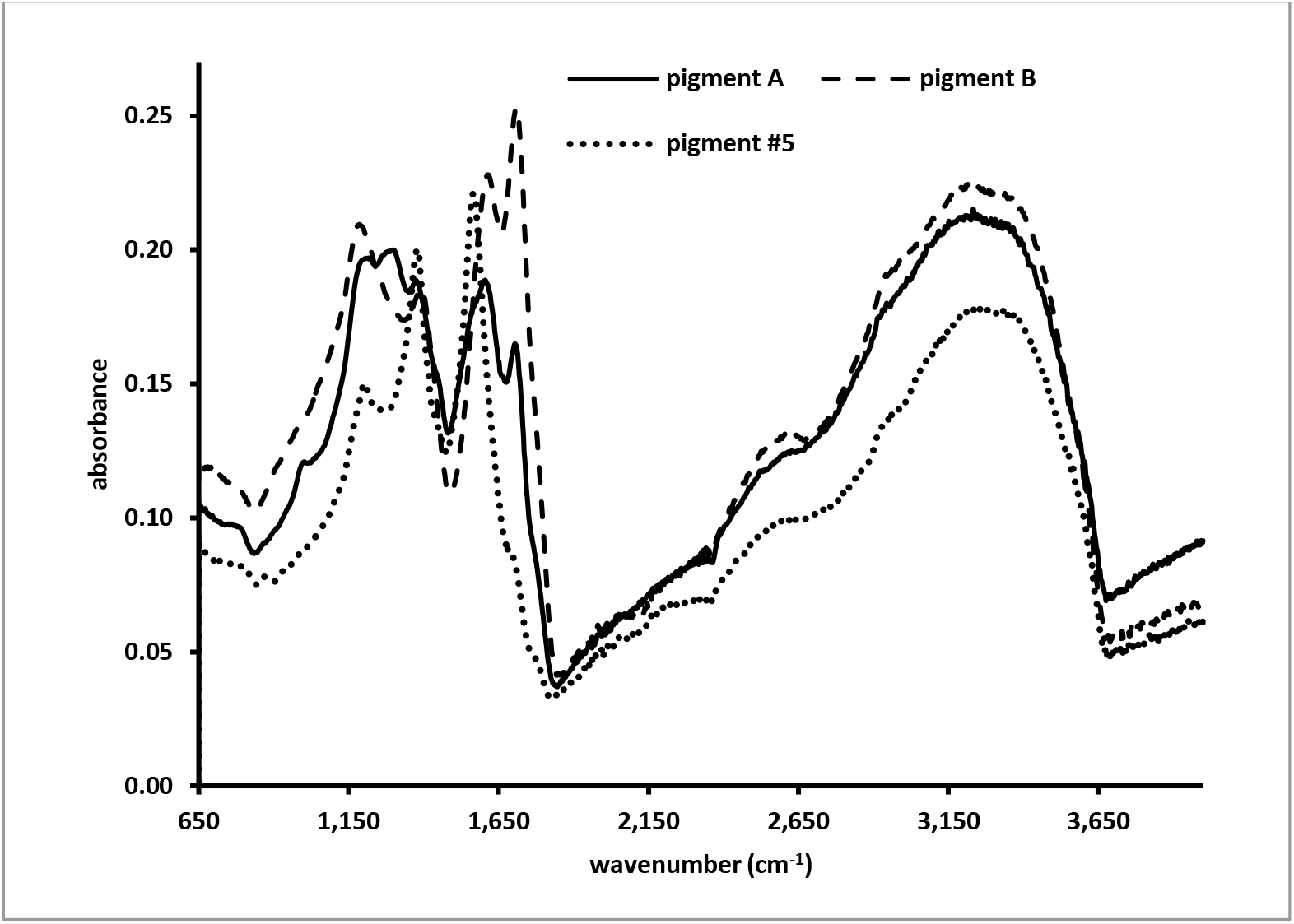
FT-IR spectra of pigments A and B, prepared as discussed in this report, and of pigment #5, prepared as discussed elsewhere[7].

Similarities and notable differences between the spectra can be observed. All three pigments exhibit absorbance peaks at wavenumbers of about 1,200cm^-1^ (associated with C-O stretching), 1,380 cm^-1^ (associated with phenolic OH-bending) and 1,600cm^-1^ (associated with aromatic C=C bending vibrations).[11] Pigment A exhibits an additional peak at a wavenumber of about 1,300 cm^-1^, the significance of which is currently unclear. Pigments A and B exhibit a sharp peak at wavenumber of about 1.705 cm^-1^ that is not present in the spectrum of pigment #5. In addition, the sharp peak at about 1.705 cm^-1^ is more prominent in the spectrum of pigment B compared to the spectrum of pigment A. We attribute this new peak to the presence of a carbonyl entity (C=O); either from a ketone or a carboxylic acid functionality.[12] This may reflect a change due to a chemical reaction whereby a phenolic functional group was converted into a ketone functional group or whereby an aromatic unit was opened and two carboxylic acids were generated into the structure. At this stage, no distinction between these two possibilities can be made. The C=O entity of aromatic carboxylic acid groups should exhibit IR absorbances at a wavenumber of about 1,750 cm^-1^.[13] However, hydrogen bonding between carboxylic acids and nearby phenolic functional groups will lead to a shift of this IR absorbance band to much lower wavenumbers. [13] The IR absorbance of C=O entities belonging to unsaturated, endocyclic ketones is predicted to appear at wavenumbers similar to the ones of hydrogen-bonded carboxylic acid groups.[13]

## 4. Discussion

The small scale experiments indicated that in the presence of Fe^2+^ and H_2_O_2_, HGA is readily converted into a darkly colored pigment. However, in the presence of the highest concentrations of H_2_O_2_ tested, the pigment’s color became lighter to the point that the solution was colorless after overnight reaction (see Fig. 1). This pattern of loss-of-colors was independent of the concentration of Fe^2+^ or the type of cation (Fe^2+^ or Cu^2+^) present. However, the results presented in Fig. 3 do suggest that the ratio of HGA and H_2_O_2_ concentrations present in the reaction mixtures may be the determining factor for the observed loss-of-color pattern. The possibility that a more intense oxidation of HGA could result in a lesser-visible pigment material was confirmed by the large scale reactions that were set up. The SEC analyses of the pigments generated through air-mediated[7] or H_2_O_2_-mediated oxidation (this report) of HGA indicate that the hydrodynamic volume (related to the molecular mass) of the pigments is not substantially different. On the other hand, a comparison of the FT-IR spectra of the pigments (Fig. 6) does suggest that H_2_O_2_-mediated oxidation leads to a chemically distinct pigment compared to air-mediated oxidation of HGA.

Our observations may have implications for the study of AKU and the associated ochronosis. HGA, at levels present in AKU patients, has been shown to be cytotoxic, particularly towards chondrocytes.[14-16] Thus, the oxidation and polymerization of HGA into a MN-like pigment could be considered a defensive mechanism to remove the excess HGA. The deposit of the HGA-based pigment leads to ochronosis, but three main questions regarding ochronosis remain: 1) how is the pigment synthesized, 2) why does it appear only in a later stage of life and 3) why is it sporadic, rather than blanket in its appearance?[6] This report suggests that darkly-colored, MN-like pigments can readily be generated from HGA in a non-enzymatic fashion due to the presence of ROS generated by Fe^2+^ and H_2_O_2_. In addition, our results suggests that, under more intensive oxidizing conditions, MN-like molecules could be generated from HGA that are much lighter in color; even to the point of being invisible. Thus, the absence of any visible pigment, as for younger AKU patients, would not necessarily mean the absence of HGA-derived, MN-like molecules. The detection of these non-visible pigments adds weight to the case for the use of nitisinone in AKU patients, given the efficacy of it to reduce HGA levels to almost baseline and removing the potential to generate any pigment; regardless of the mechanism of pigment formation.[17, 18] It is interesting to compare the relationship between AKU and ochronosis and other conditions or diseases that involve changes in pigmentation: vitiligo or greying of hair.[19, 20] For both, an increased level of H_2_O_2_ has been discussed a potential contribution factor to the changes in pigmentation. [9, 10] In this respect is worth noting that HGA can auto-oxidize and produce ROS like H_2_O_2_.[21] Although the effect of age on pigmentation in AKU (from colorless to darkly colored) is the reverse as that for aging hair (from darkly colored to lighter colored), we can expand our comparison between ochronosis and greying of hair by noting that, with regard to the biochemical and physiological changes in hair, the third decade is considered to be a pivot point. [20] Both pathologies are also associated with a compromised ability to defend against ROS.[22-24] The pigmentation associated with AKU and ochronosis is derived from HGA, while other precursors, cateholamines or DOPA, form the basis for the pigmentation associated with vitiligo or hair color. We have observed that in addition to HGA, other catecholic precursors can generate light-colored MN-like materials when oxidized in the presence of high concentrations of H_2_O_2_ and the results of these observations will be presented elsewhere.

It is also worth considering the *in vivo* location of the most severe pigmentation in AKU patients, the articular cartilage. This tissue is usually avascular and aneural, but numerous disease processes or trauma bring about changes in cartilage. It may be these micro-trauma events that instigate the pigmentation process due to the generation of additional ROS or the through the increased availability of ions like Fe^2+^ as part of the repair response mechanism. These micro-changes in the cartilage of AKU patients are well characterized as pericellular pigmentation and are clearly the initiation point in the process of generating visible ochronosis.[25, 26]

In general, we present evidence that the H_2_O_2_-mediated oxidation of HGA can lead to a MN-like material that is chemically distinct from pigments generated through air-oxidation. In addition, we provide evidence that intense oxidation by H_2_O_2_ can lead to MN-like materials with reduced absorbance in the visible range of the electromagnetic spectrum. The prospect of the existence of HGA-derived, non-ochronotic pigments in addition to the well-established ochronotic pigments, could have implications in the study and therapy of AKU/ochronosis as the absence of pigmentation observed in younger AKU patients may not mean the lack of MN-like materials.

## Acknowledgements

Dr. Vercruysse was in part supported by the Institute for Food, Agricultural and Environmental Research at Tennessee State University. Dr Taylor was in part supported by the Rosetrees Trust.

## References

[1] J.B. Mistry, M. Bukhari, A.M. Taylor, Alkaptonuria, Rare Dis 1 (2013) e27475.

[2] A. Zatkova, An update on molecular genetics of Alkaptonuria (AKU), J Inherit Metab Dis 34(6) (2011) 1127–36.

[3] D. Braconi, L. Millucci, G. Bernardini, A. Santucci, Oxidative stress and mechanisms of ochronosis in alkaptonuria, Free Radical Biology and Medicine 88, Part A (2015) 70–80.

[4] J.M. Keller, W. Macaulay, O.A. Nercessian, I.A. Jaffe, New developments in ochronosis: review of the literature, Rheumatol Int 25(2) (2005) 81–5.

[5] N.B. Roberts, S.A. Curtis, A.M. Milan, L.R. Ranganath, The Pigment in Alkaptonuria Relationship to Melanin and Other Coloured Substances: A Review of Metabolism, Composition and Chemical Analysis, JIMD Rep 24 (2015) 51–66.

[6] A.M. Taylor, V. Kammath, A. Bleakley, Tyrosinase, could it be a missing link in ochronosis in alkaptonuria?, Med Hypotheses 91 (2016) 77–80.

[7] A.M. Taylor, K.P. Vercruysse, Analysis of Melanin-like Pigment Synthesized from Homogentisic Acid, with or without Tyrosine, and Its Implications in Alkaptonuria, Springer Berlin Heidelberg, Berlin, Heidelberg, pp. 1–7.

[8] Y.T. Konttinen, J. Mandelin, T.-F. Li, J. Salo, J. Lassus, M. Liljeström, M. Hukkanen, M. Takagi, I. Virtanen, S. Santavirta, Acidic cysteine endoproteinase cathepsin K in the degeneration of the superficial articular hyaline cartilage in osteoarthritis, Arthritis & Rheumatism 46(4) (2002) 953–960.

[9] K.U. Schallreuter, J. Moore, J.M. Wood, W.D. Beazley, D.C. Gaze, D.J. Tobin, H.S. Marshall, A. Panske, E. Panzig, N.A. Hibberts, In vivo and in vitro evidence for hydrogen peroxide (H_2_O_2_) accumulation in the epidermis of patients with vitiligo and its successful removal by a UVB-activated pseudocatalase, J Investig Dermatol Symp Proc 4(1) (1999) 91–6.

[10] J.M. Wood, H. Decker, H. Hartmann, B. Chavan, H. Rokos, J.D. Spencer, S. Hasse, M.J. Thornton, M. Shalbaf, R. Paus, K.U. Schallreuter, Senile hair graying: H_2_O_2_-mediated oxidative stress affects human hair color by blunting methionine sulfoxide repair, FASEB J 23(7) (2009) 2065–75.

[11] C.E. Turick, L.S. Tisa, F. Caccavo, Jr., Melanin production and use as a soluble electron shuttle for Fe(III) oxide reduction and as a terminal electron acceptor by Shewanella algae BrY, Appl Environ Microbiol 68(5) (2002) 2436–44.

[12] J. Coates, Interpretation of Infrared Spectra, A Practical Approach, Encyclopedia of Analytical Chemistry, John Wiley & Sons, Ltd 2006.

[13] E. Fuente, J.A. Menéndez, M.A. Díez, D. Suárez, M.A. Montes-Morán, Infrared Spectroscopy of Carbon Materials: A Quantum Chemical Study of Model Compounds, The Journal of Physical Chemistry B 107(26) (2003) 6350–6359.

[14] D. Braconi, G. Bernardini, C. Bianchini, M. Laschi, L. Millucci, L. Amato, L. Tinti, T. Serchi, F. Chellini, A. Spreafico, A. Santucci, Biochemical and proteomic characterization of alkaptonuric chondrocytes, J Cell Physiol 227(9) (2012) 3333–43.

[15] C.J. Kirkpatrick, W. Mohr, W. Mutschler, Experimental studies on the pathogenesis of ochronotic arthropathy. The effects of homogentisic acid on adult and fetal articular chondrocyte morphology, proliferative capacity and synthesis of proteoglycans in vitro, Virchows Arch B Cell Pathol Incl Mol Pathol 47(4) (1984) 347–60.

[16] J.B. Mistry, D.J. Jackson, M. Bukhari, A.M. Taylor, A role for interleukins in ochronosis in a chondrocyte in vitro model of alkaptonuria, Clin Rheumatol 35(7) (2016) 1849–56.

[17] A.M. Milan, A. Hughes, A.S. Davison, J.M. Devine, J.L. Usher, S. Curtis, M. Khedr, J.A. Gallagher, L. Ranganath, ANNALS EXPRESS: The effect of nitisinone on homogentisic acid and tyrosine: A two-year survey of patients attending the National Alkaptonuria Centre, Liverpool, Ann Clin Biochem (2017) 4563217691065.

[18] L.R. Ranganath, A.M. Milan, A.T. Hughes, J.J. Dutton, R. Fitzgerald, M.C. Briggs, H. Bygott, E.E. Psarelli, T.F. Cox, J.A. Gallagher, J.C. Jarvis, C. van Kan, A.K. Hall, D. Laan, B. Olsson, J. Szamosi, M. Rudebeck, T. Kullenberg, A. Cronlund, L. Svensson, C. Junestrand, H. Ayoob, O.G. Timmis, N. Sireau, K.H. Le Quan Sang, F. Genovese, D. Braconi, A. Santucci, M. Nemethova, A. Zatkova, J. McCaffrey, P. Christensen, G. Ross, R. Imrich, J. Rovensky, Suitability Of Nitisinone In Alkaptonuria 1 (SONIA 1): an international, multicentre, randomised, open-label, no-treatment controlled, parallel-group, dose-response study to investigate the effect of once daily nitisinone on 24-h urinary homogentisic acid excretion in patients with alkaptonuria after 4 weeks of treatment, Ann Rheum Dis 75(2) (2016) 362–7.

[19] K.U. Schallreuter, P. Bahadoran, M. Picardo, A. Slominski, Y.E. Elassiuty, E.H. Kemp, C. Giachino, J.B. Liu, R.M. Luiten, T. Lambe, I.C. Le Poole, I. Dammak, H. Onay, M.A. Zmijewski, M.L. Dell’Anna, M.P. Zeegers, R.J. Cornall, R. Paus, J.P. Ortonne, W. Westerhof, Vitiligo pathogenesis: autoimmune disease, genetic defect, excessive reactive oxygen species, calcium imbalance, or what else?, Exp Dermatol 17(2) (2008) 139–40; discussion 141-60.

[20] D.J. Tobin, Age-related hair pigment loss, Curr Probl Dermatol 47 (2015) 128–38.

[21] J.P. Martin, Jr., B. Batkoff, Homogentisic acid autoxidation and oxygen radical generation: implications for the etiology of alkaptonuric arthritis, Free Radic Biol Med 3(4) (1987) 241–50.

[22] R.F. Loeser, Aging and osteoarthritis: the role of chondrocyte senescence and aging changes in the cartilage matrix, Osteoarthritis Cartilage 17(8) (2009) 971–9.

[23] Y. Shi, L.F. Luo, X.M. Liu, Q. Zhou, S.Z. Xu, T.C. Lei, Premature graying as a consequence of compromised antioxidant activity in hair bulb melanocytes and their precursors, PLoS One 9(4) (2014) e93589.

[24] A.M. Taylor, M.F. Hsueh, L.R. Ranganath, J.A. Gallagher, J.P. Dillon, J.L. Huebner, J.B. Catterall, V.B. Kraus, Cartilage biomarkers in the osteoarthropathy of alkaptonuria reveal low turnover and accelerated ageing, Rheumatology (Oxford) 56(1) (2017) 156–164.

[25] A.J. Preston, C.M. Keenan, H. Sutherland, P.J. Wilson, B. Wlodarski, A.M. Taylor, D.P. Williams, L.R. Ranganath, J.A. Gallagher, J.C. Jarvis, Ochronotic osteoarthropathy in a mouse model of alkaptonuria, and its inhibition by nitisinone, Ann Rheum Dis 73(1) (2014) 284–9.

[26] A.M. Taylor, A. Boyde, P.J. Wilson, J.C. Jarvis, J.S. Davidson, J.A. Hunt, L.R. Ranganath, J.A. Gallagher, The role of calcified cartilage and subchondral bone in the initiation and progression of ochronotic arthropathy in alkaptonuria, Arthritis Rheum 63(12) (2011) 3887–96.

